## Supplementary Information SI1 for "Modelling protein complexes with crosslinking mass spectrometry and deep learning"

**Supplementary Information SI1: The FpaA (YlaN) protein acts as an antirepressor to the ferric uptake regulator Fur in *Bacillus subtilis***

All living organisms require the presence of micronutrients to survive among them iron. As many other essential metal ions, iron is not only required for life, but its accumulation can also have toxic effects.

Since iron is essential but also toxic, intracellular iron levels need to be tightly regulated. This regulation is mediated by Fur, the ferric uptake regulator which is conserved in many archaea and most bacteria. Fur belongs to the ferric uptake regulator family which includes Zur, the major regulator of zinc homeostasis and PerR which responds to peroxide stress. Fur regulates the expression of at about 60 genes in *Bacillus subtilis* (1, 2). Fur almost exclusively represses the expression of its target genes which are mostly iron uptake and siderophore synthesis systems (1). Fur consists of an N-terminal DNA-binding domain and a C-terminal domain which is thought to bind ferrous iron as a cofactor and which mediates dimerization (3, 4). It has long been assumed that Fur binds ferrous iron as a cofactor but only recently it could be shown that Fur actually reversibly binds an iron sulfur cluster in *Escherichia coli* (5).

Even though it could not be shown, it is widely assumed that *B. subtilis* Fur senses the intracellular iron concentration by binding ferrous iron ($Fe^{2+}$) (6, 7). With decreasing extracellular iron concentrations the Fur regulon is derepressed in three waves in *B. subtilis* (2). First, iron uptake systems for elemental iron, ferric citrate and petrobactin are expressed. Sequentially, the synthesis of the siderophore bacillibactin and uptake systems for bacillibactin and the hydroxamate siderophores to scavenge iron occurs before the iron-sparing response is initiated to inhibit the translation of iron binding proteins.

Despite the absence of direct evidence for the binding of iron to Fur, there has been only little research regarding possible other mechanisms that regulate Fur. However, recent studies revealed that there are proteins which modulate the activity of Fur in different bacteria (8). In *Salmonella enterica*, the $EIIA^{Ntr}$ protein of the non-canonical phosphotransferase system regulates the expression of iron uptake genes via Fur by a direct protein-protein interaction which results in the release of Fur from its DNA binding sites (9). Another but similar mechanism was recently found in uropathogenic *E. coli*, where the proteins YdiV and SlyD cooperatively bind Fur and reduce its DNA binding (10). These results indicate that the regulation of Fur, which was assumed to be exclusively dependend on ferrous iron, actually involves Fur antagonists that might be more common then previously anticipated.

We are interested in the functional characterization of unknown or poorly studied proteins in the model bacterium *B. subtilis*. The YlaN protein a highly abundant but only poorly studied protein (11, 12). Moreover, the *ylaN* gene is essential under standard growth conditions (13, 14). Both the very high expression and the essentiality suggest that the YlaN protein plays a very important role in the cell. Recently it was shown that t he *ylaN* gene becomes dispensable when ferric iron is added to the growth medium (15). This provides a strong indication that the YlaN protein is involved in the control of iron homeostasis. Moreover, recent *in vivo* crosslinking data revealed an intriguing interaction between YlaN and Fur which indicates that YlaN might be another Fur antagonist and thus exert its role in iron homeostasis via Fur (16, 17).

Here, we have studied the role of YlaN in the control of iron homeostasis in *B. subtilis*. We confirm the direct protein-protein interaction between Fur and YlaN, and show that Fur is unable to bind its target DNA in the presence of YlaN. Thus, YlaN acts as Fur protein antagonist which we rename Fpa.

**Fpa becomes dispensable in the absence of Fur.**

It has been shown that the essential *fpa* (previously *ylaN*) gene becomes dispensable if ferric iron is added to the growth medium (15). Based on the interaction between Fpa and the Fur regulator protein, we hypothesized that Fpa might antagonize Fur to allow the expression of iron uptake systems at low iron concentrations. If this were true, the deletion of the *fpa* gene under standard conditions might be toxic as a result of continued repression of the genes for iron uptake by Fur. To test this idea, we attempted the deletion of *fpa* in the *fur* mutant GP879. As a control, we used the isogenic wild type strain *B. subtilis* 168. In agreement with the published data, *fpa* could not be deleted under standard conditions or if the plates were supplemented with ferrous iron. However, the deletion was possible if the medium was supplemented with ferric iron ($Fe^{3+}$). In contrast, the *fpa* gene could be deleted under all three conditions in the *fur* mutant strain GP879. These results confirm the conditional essentiality of Fpa depending on the iron supply and they suggest that Fpa might be needed to prevent some harmful activity of Fur under conditions of iron limitation.

**Fpa is quasi-essential under standard growth conditions.**

When the concept of essentiality was introduced, a gene was regarded essential if it could not be activated under standard growth conditions for the organism, *i. e*. on LB medium supplemented with glucose at 37°C for *B. subtilis* (14). Today, we know that several essential genes can in fact be deleted under standard conditions but that the mutant strains immediately acquire suppressor mutations that help to overcome the growth-limiting problem. Such genes are called quasi-essential as they remain essential under standard conditions in the genomic context of the standard wild type strain.

The possibility to delete *fpa* from a *fur* mutant suggested that *fpa* might also be quasi-essential. Thus, we made use of the *fpa* mutant GP3324 that had been isolated in the presence of ferric iron as described above. The strain was cultivated on complex medium under iron-limiting condition. As expected, we observed the development of few individual colonies that most likely resulted from the acquisition of suppressor mutations. We re-isolated eight independent colonies that had appeared in the absence of added iron or in the presence of ferrous iron each, and sequenced their *fur* alleles as we already knew that *fur* mutants tolerate the inactivation of *fpa*. Of 15 individual mutants, all had single point mutations in the Fur coding sequence that resulted in amino acid substitutions in the Fur protein. These mutations all occurred in the N-terminal part of the protein which is required for DNA binding indicating that the mutated Fur versions are impaired in DNA binding (3). Several of the substitutions were observed in multiple mutants, both from selection without added iron or in the presence of ferrous iron (see Fig. 1A).

These results demonstrate that *fpa* is a novel quasi-essential gene. Under standard conditions on complex medium, the deletion of *fpa* causes the immediate acquisition of suppressor mutations that interfere with Fur activity. Moreover, these results support the idea that the Fur repressor becomes toxic for the cells in the absence of Fpa and iron, i.e. under conditions that require expression of the iron uptake genes that are all members of the Fur regulon.

**Fpa affects the activity of the Fur-controlled *dhbA* promoter.**

The *dhbACEBF-ybdZ* operon encodes the enzymes for the synthesis of the siderophore bacillibactin (18). It is one of the Fur-controlled operons, and it is induced as a part of the second wave of Fur-controlled genes that are expressed upon iron limitation (2). We used the activity of the *dhbA* promoter as an indicator of Fur activity. For this purpose, we fused the *dhbA* promoter region to a promoterless *lacZ* gene encoding β-galactosidase and integrated this *dhbA-lacZ* fusion into the *B. subtilis* genome of the wild type strain and the isogenic Δ*fur* mutant GP879. The resulting strains, GP3331 and GP3356, respectively, carrying the *dhbA-lacZ* fusion were cultivated in CSE-Glc minimal medium supplemented with 0.1 µM and 500 µM of ferrous and ferric iron. As shown in Fig. 2, a strong promoter activity was determined at 0.1 µM iron, irrespective of the redox state of the iron ions. Expression was substantially reduced at the increased iron concentration in the wild type strain. For ferrous iron, we observed an about twofold decrease of expression, whereas a twelvefold decrease was detected in the case of ferric iron. This result corresponds well to the observation that ferrous iron is less bioavailable for the bacteria, so that it causes only a weak repression of *dhbA* expression. In the *fur* mutant GP3356, we observed high β-galactosidase under all tested conditions. The strong repression of *dhbA* expression in response to ferrous iron, and the dependence of repression on a functional Fur protein is in excellent agreement with published data (19). We also tested the expression of the *dhbA-lacZ* fusion in the wild type strain GP3331 in complex medium (LB) in the presence and absence of 500 µM ferrous or ferric iron. In this case, the activity was already rather low in the absence of added iron (45 unit per mg of protein as compared to 700 ... 900 units in minimal medium in the presence of 0.1 µM of iron), probably due to the presence of iron in the complex medium which is likely to cause a basal repression. However, the expression was fivefold repressed if one of the iron ions was added (see Fig. 3A for $Fe^{3+}$, data not shown for $Fe^{2+}$). For further experiments, we used LB medium and ferric iron.

To test the role of Fpa in the Fur-mediated regulation of the *dhbA* promoter, we constructed additional strains carrying the *dhbA-lacZ* fusion in a *fpa* single and *fpa fur* double mutant. The resulting strains were GP3366 and GP3361, respectively. As shown in Fig. 3A, expression in the *fur* mutant was strongly enhanced as compared to the wild type background and independent from the iron concentration. The rather weak expression in the wild type strain in the absence of added iron is in good agreement with the conclusion that this medium already contains some iron. The loss of Fpa in addition to Fur in GP3361 had no effect on the *dhbA* promoter activity. Finally, we tested *dhbA* promoter activity in the *fpa* mutant (GP3366). For this strain, only background activities were observed under both conditions. We conclude that Fpa exerts its role via Fur, as already suggested as a result from the suppression analysis. The lack of promoter activity in the *fpa* single mutant is in excellent agreement with the above conclusion that the Fur-controlled genes for iron uptake might be completely repressed in the absence of Fpa thus resulting in the essentiality of Fpa in standard LB medium.

The deletion of *fpa* resulted in complete repression of the *dhbA* promoter, likely due to the inability to release Fur from its target DNA in the absence of Fpa. Given the observed physical interaction between the two proteins, it is tempting to speculate that the overexpression of Fpa might cause the release of Fur from its target sites and thus induction of *dhbA* expression even in the presense of iron. To test this hypothesis, we put the *fpa* gene under the control of the strong *degQ*$^{hy}$ promoter in the expression plasmid pBQ200. The resulting plasmid was pGP3897. This plasmid as well as the empty vector pBQ200 were then introduced into strain GP3331 that harbors the *dhbA-lacZ* fusion. Again, the strains were grown in LB medium in the presence or absence of 500 µM ferric citrate. As observed before for the wild type strain, we found a fivefold repression of *dhbA* expression in the presence of the empty vector (see Fig. 3B). In contrast, expression was strongly increased in the presence of pGP3897 when Fpa was overproduced. We even observed a substantial expression in this strain if 500 µM ferric citrate were present. This observation indicates that the overexpression of Fpa counteracts the repressing effect caused by Fur and strongly suggests that Fpa acts as a *bona fide* antagonist of Fur.

**Physical interaction between Fur and Fpa.** Our previous proteome-wide interaction studies with *B. subtillis* detected an interaction between Fur and Fpa (17). If Fpa acts as an antagonist to Fur, it seems most likely that this activity is achieved by the physical interaction between the two proteins. To confirm this interaction, we decided to verify the interaction *in vivo* in *B. subtilis* by co-precipitation. For this purpose, we constructed a strain that expressed Fur carrying a C-terminal FLAG tag for immunological detection (GP3367). This strain was then transformed either with the empty vector pGP382 (20) or with plasmid pGP3867 for the expression of Fpa fused to a C-terminal Strep-tag for affinity chromatography. Both strains were cultivated in CSE-Glc minimal medium with 0.1 or 500 µM ferric citrate. The protein extracts were then passed over a StrepTactin column to bind the Fpa-Strep, washed and Strep-tagged proteins with their potential interaction partners were eluted. Two proteins, PycA and AccB, were eluted from the StrepTactin column for both the empty vector control and the strain expressing Fpa-Strep. These proteins contain a biotin cofactor that causes binding to the matrix and are good indicators that the experimental setup was suitable. Upon expression of Fpa-Strep, we observed copurification of a protein of about 20 kDa which corresponds to FLAG-tagged Fur. The identity of the band was confirmed by a Western blot using antibodied directed against the FLAG tag (see Fig. 4). The Fur protein was copurified with Fpa both at high and low iron concentrations. Again, the low iron concentration used in this experiment may still be sufficient to allow the interaction between the two proteins.

**Fpa prevents the DNA-binding activity of Fur.** All our experiments support our initial hypothesis that Fpa can bind Fur and interfere with the repression of target genes by Fur. To get direct evidence for this, we performed DNA binding assays with the *dhbA* promoter region and purified Fur protein. As shown in Fig. 5, the DNA fragment was retarded in the presence of Fur, indicating binding of Fur to its target DNA. In contrast, no shift was observed with the purified Fpa protein. The addition of both Fur and Fpa to the DNA did not result in DNA binding. This observation is in excellent agreement with the idea that Fpa might prevent Fur from binding to DNA. To ensure that the effect of Fpa addition to the DNA and Fur is specific, we performed a control experiment in which we used the PtsH protein of the phosphotransferase system as the seond protein in the assay. As Fpa, PtsH is a small acidic protein. The results observed with the promoter DNA and Fur were as described above. Similarly, the PtsH protein did not interact with the DNA. However, the presence of PtsH in addition to the promoter fragment and Fur did not prevent the formation of the Fur-DNA complex indicating that PtsH is unable to interfere with the DNA binding activity of Fur (Fig. 5). Taken together, our results demonstrate that Fpa specifically interacts with Fur to prevent it from binding to its target DNA sequences and thus to allow the expression of genes that are under negative control by Fur.

### Conclusion

The Fpa (YlaN) protein belongs to a small group of so far unknown proteins that are strongly expressed under essentially all conditions in *B. subtilis* (12). Of those about 40 proteins, Fpa is the only that is essential under standard laboratory growth conditions (13). The data presented in this study identify the so far unknown protein Fpa as a *bona fide* antirepressor of the Fur regulator. The interaction between the two proteins interferes with the binding of Fur to its DNA targets and thus results in the expression of the iron uptake systems which are all subject to Fur-mediated transcription repression.

Fpa perceives the primary signal of the system, the intracellular iron concentration in the form of ferrous iron (21). In this form, the protein does not interact with Fur, and Fur can bind to its DNA target sites to repress the expression of genes for iron uptake systems and to activate expression of an iron exporter gene. In contrast, under conditions of iron limitation, apo-Fpa can bind to Fur, and thus break open the Fur dimer, resulting in release of Fur from its target sites as shown in this work. This interaction allows the expression of Fur-controlled genes for iron uptake systems if iron gets scarce, and is thus a prerequisite for the growth of *B. subtilis* under conditions of iron depletion. This explains why the *fpa* gene is essential for *B. subtilis* under standard conditions. Since iron limitation is the rule rather than the exception for bacteria that live under aerobic conditions, mechanisms that allow the effective induction of genes for iron acquisition are of key importance.

Fur-mediated control of iron homeostasis is widespread in both Gram- negative and Gram-positive bacteria. For long time, it was assumed that the Fur protein directly responds to the presence of ferrous iron (22). However, despite intensive research there is no clear support for this idea in the published data body and the direct sensing of iron by Fur has remained controversial (23, 24). In contrast, there is clear evidence that zinc ions act as cofactor for the regulator of zinc homeostasis Zur, another Fur-type regulator (25). Only more recent studies with Gram-negative bacteria and the data presented here suggest that Fur may be controlled by regulatory protein-protein interactions in many bacteria (8).

An analysis of the phylogenetic distribution of Fpa reveals that the protein is present exclusively in the Bacilli subgroup of the Firmicutes (26). In this class, Fpa is present in most species with the exception of the lactic acid bacteria and few other species. Interestingly, most bacteria that possess Fur family transcription factors, encode multiple, typically three of these proteins, Fur, PerR, and Zur. Those bacteria of the Bacilli class that lack Fpa, do also lack the Fur protein. Most of them have PerR and Zur, with the notable exception of *Lactobacillus acidophilus* and *Streptococcus pneumoniae* that completely lack Fur type regulators. In the genus *Jeotgalibacillus*, one species, *J. malaysiensis* possesses both Fur and Fpa, whereas *J. donkookensis* encodes neither of the two proteins. There are only two bacteria among the Bacilli that seem to possess Fur but not Fpa, *Aneurinibacillus soli* CB4 and *Tumebacillus avium* AR23208. This might result from issues with the genome sequences, or these bacteria have evolved specific strategies to control Fur activity.

The two-faced role of iron as an important player in cellular energy metabolism on one hand and its toxicity on the other make the presence of effective systems to control iron homeostasis critical for bacterial life. As a result of Fur/Fpa-dependent regulation, the genes for iron uptake are not expressed if the metal is already abundant in the cell. The antirepressor activity of Fpa allows expression of iron uptake genes as soon as iron gets limiting. This is a strategic decision of the cell as it determines which protein of the iron homeostasis system will be present or not in the future.

The discovery of the activity of Fpa as an antirepressor to Fur in *B. subtilis* is important for our better understanding of the physiology of this important model organism (27). *B. subtilis* is the model organism for a large group of Gram-positive bacteria, that includes many important pathogens such as *S. aureus*, *Listeria monocytogenes*, or *Bacillus anthracis* that also possess the Fur/Fpa couple. Iron is the growth-limiting factor for most pathogenic bacteria in the human body. Accordingly, the investigation of the control of iron homeostasis is also very important to better understand the processes of infection and disease and to develop novel treatments. Since Fpa is essential under conditions of iron limitation not only in *B. subtilis* but also in *S. aureus* (21), it might be an attractive novel target for drug development.

### SUPPLEMENTAL METHODS

**Genome sequencing.** To identify the mutations in the suppressor mutant strain GP3368 (see Table 1), the genomic DNA was subjected to whole-genome sequencing. Concentration and purity of the isolated DNA was first checked with a Nanodrop ND-1000 (PeqLab Erlangen, Germany), and the precise concentration was determined using the Qubit® dsDNA HS Assay Kit as recommended by the manufacturer (Life Technologies GmbH, Darmstadt, Germany). Illumina shotgun libraries were prepared using the Nextera XT DNA Sample Preparation Kit and subsequently sequenced on a MiSeq system with the reagent kit v3 with 600 cycles (Illumina, San Diego, CA, USA) as recommended by the manufacturer. The reads were mapped on the reference genome of *B. subtilis* 168 (GenBank accession number: NC_000964) (28). Mapping of the reads was performed using the Geneious software package (Biomatters Ltd., New Zealand) (29). Frequently occurring hitchhiker mutations (30) and silent mutations were omitted from the screen. The resulting genome sequence was compared to that of our in-house wild type strain. Single nucleotide polymorphisms were considered as significant when the total coverage depth exceeded 25 reads with a variant frequency of ≥90%. All identified mutations were verified by PCR amplification and Sanger sequencing.

***In vivo* detection of protein-protein interactions.** For the expression of Fpa carrying a C-terminal Strep-tag in *B. subtilis*, we used plasmid pGP3867. This plasmid was obtained by cloning the *fpa* gene between the BamHI and SalI sites of the expression vector pGP382 (20). To add a FLAG tag epitope to the Fur protein, we constructed plasmid pGP3899 by cloning the *fur* gene between the BamHI and HindIII sites of pGP1331 (31). Transformation of the wild type strain 168 with pGP3899 yielded strain GP3367. To verify the interaction between Fur and Fpa *in vivo*, cultures of *B. subtilis* GP3367 (Fur-FLAG) containing pGP3867 (Fpa-Strep), or the empty vector control (pGP382), were cultivated in 500 ml CSE-Glc medium containing the indicated iron source until exponential growth phase was reached ($OD_{600}$ ~ 0.4-0.6). The cells were harvested immediately and stored at -20°C. The Strep-tagged protein and its potential interaction partners were then purified from crude extracts using a StrepTactin column (IBA, Göttingen, Germany) and D-desthiobiotin as the eluent. The eluted proteins were separated on an SDS gel and potential interacting partners were analyzed by staining with Colloidal Coomassie and Western blot analysis using antibodies raised against the FLAG-tag.

**Table 1.** *B. subtilis* strains.

| Strain | Genotype | Source or Reference |
|---|---|---|
| 168 | *trpC2* | Laboratory collection |
| GP879 | *trpC2* Δ*fur*::*ermC* | This study |
| GP3321 | *trpC2* Δ*fur*::*ermC* Δ*fpa*::*cat* | This study |
| GP3324 | *trpC2* Δ*fpa*::*cat* | This study |
| GP3331 | *trpC2 amyE*::($P_{dhbA}$-*lacZ aphA3*) | pGP3594 → 168 |
| GP3356 | *trpC2* Δ*fur*::*ermC amyE*::($P_{dhbA}$-*lacZ aphA3*) | GP879 → GP3331 |
| GP3361 | *trpC2* Δ*fur*::*ermC* Δ*fpa*::*cat amyE*::($P_{dhbA}$-*lacZ aphA3*) | GP3321 → GP3331 |
| GP3366 | *trpC2* Δ*fpa*::*cat amyE*::($P_{dhbA}$-*lacZ aphA3*) | GP3324 → GP3331 |
| GP3367 | *trpC2 fur*-FLAG *spc* | pGP3899 → 168 |

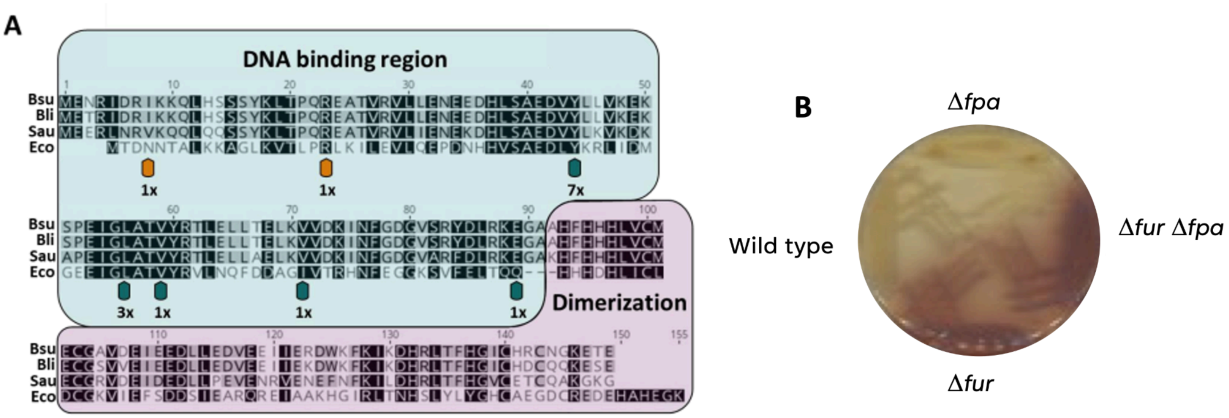


**Figure 1. Mutations in Fur upon deletion of the *fpa* gene.**

**A.** Alignment of the sequences of Fur proteins from different bacteria. The DNA-binding and dimerization domains are highlighted in light green and magenta, respectively. The positions of point mutations in the individual suppressor mutants are shown by arrows. The numbers indicate how often an individual amino acid substitution was found. The orange arrows indicate that the corresponding mutant colonies were reddish as the D*fur* mutant. Bsu, *B. subtilis*; Bli, *Bacillus licheniformis*; Sau, *S. aureus*; Eco, *E. coli*. **B.** Growth of wild type and mutant strains of *B. subtilis* on a LB agar plate containing 2.5 mM ferric iron. The deletion of the *fur* gene results in a red colony color.

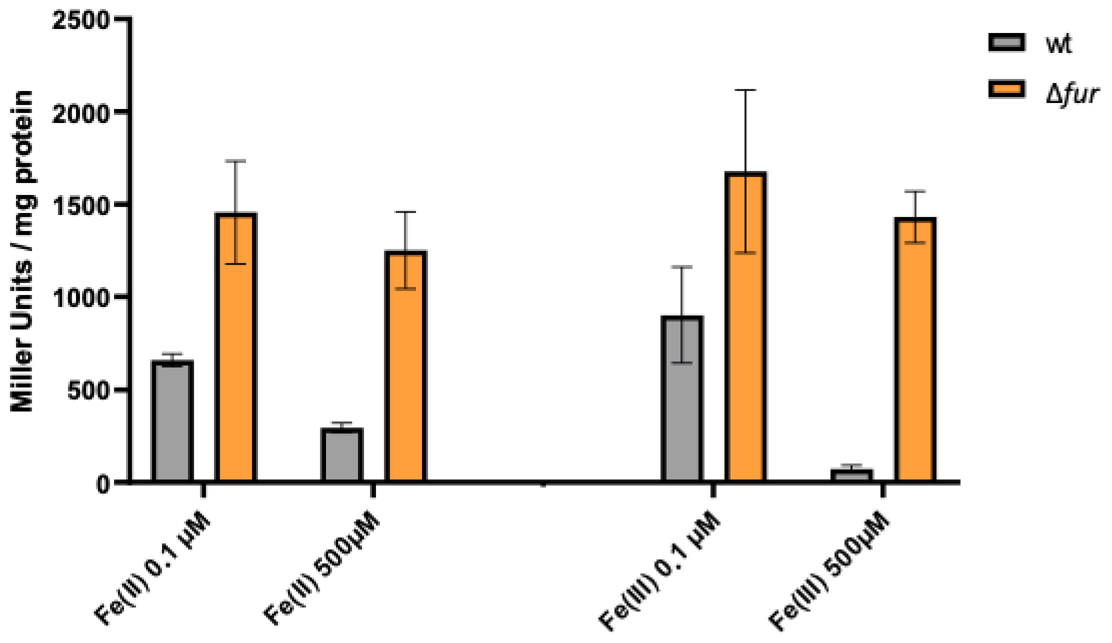


**Figure 2. Effect of iron on the activity of the *dhbA* promoter.**

Cultures of a wild type strain (GP3331) and the isogenic D*fur* mutant (GP3356) carrying a *dhbA-lacZ* fusion were grown with the indicated iron supplementation, and promoter activities were determined by quantification of b-galactosidase activities. The values are averages of three independent experiments. Standard deviations are shown.

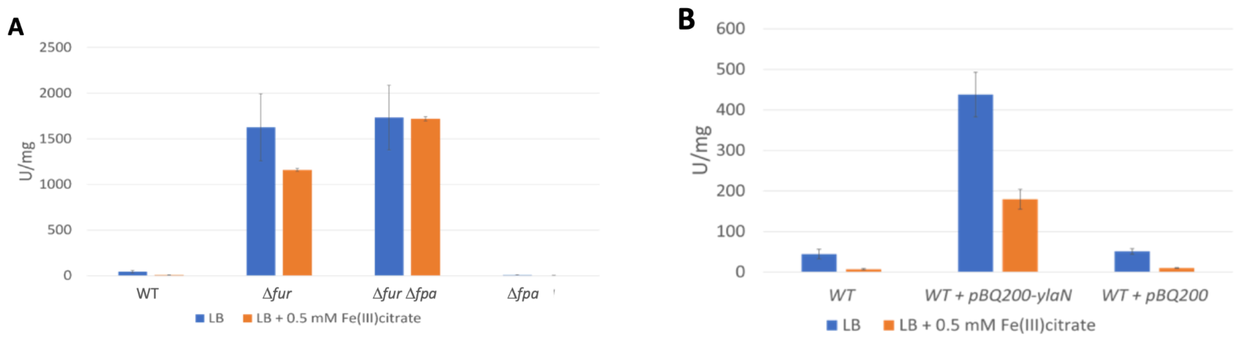


**Figure 3. The impact of Fpa on *dhbA* promoter activity.**

**A.** Effect of *fpa* deletion. Strains carrying a *dhbA-lacZ* fusion were cultivated in LB with or without added ferric citrate, and promoter activities were determined by quantification of b-galactosidase activities. The values are averages of three independent experiments. Standard deviations are shown. WT, GP3331; D*fur*, GP3356, D*fur* D*fpa*, GP3361; D*fpa*, GP3366. **B.** Effect of *fpa* overexpression. *B. subtilis* GP3331 without any plasmid (WT) and GP3331 carrying plasmid pGP3897 for *fpa* overexpression or the empty vector pBQ200 were cultivated in LB with or without added ferric citrate, and promoter activities were determined by quantification of b-galactosidase activities. The values are averages of three independent experiments. Standard deviations are shown.

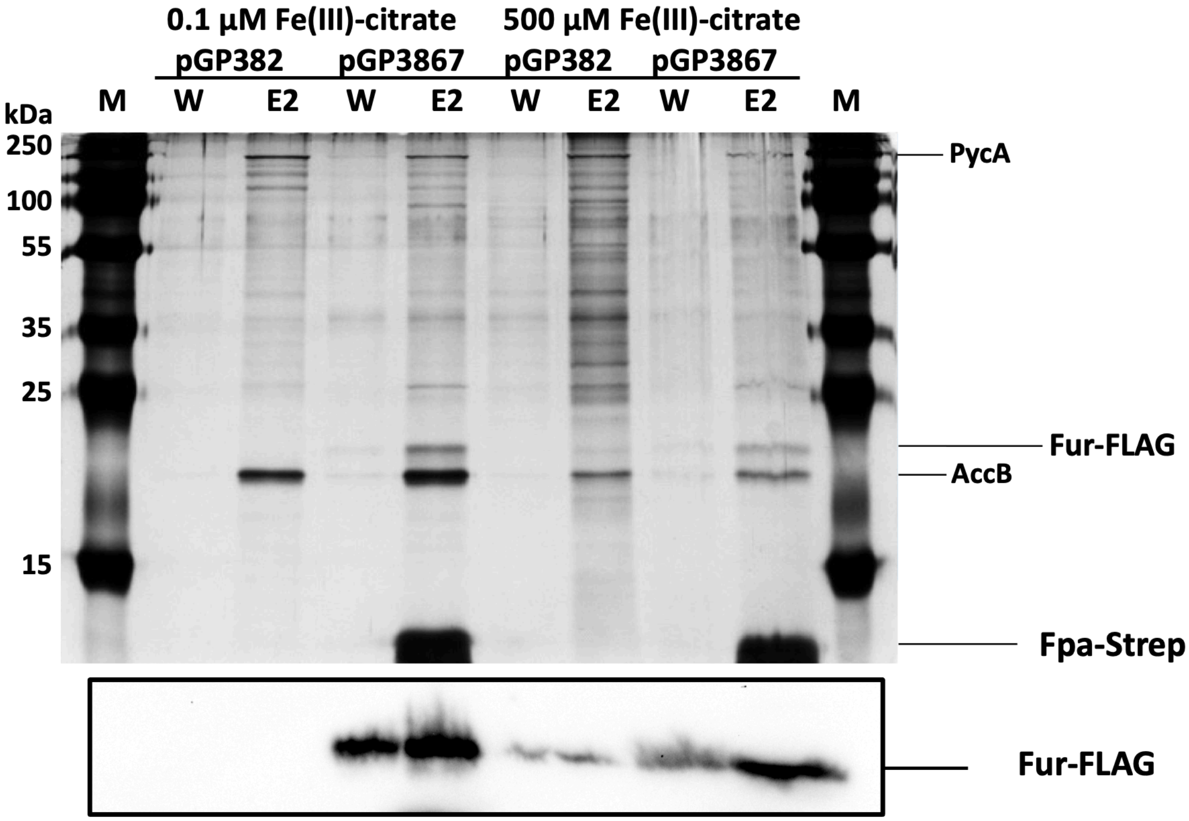


**Figure 4. Physical interaction between Fpa and Fur.**

Protein complexes isolated from *B. subtilis* GP3367 (Fur-FLAG) containing either the empty vector pGP382 or pGP3867 (Fpa-Strep). The strains were grown in CSE-Glc minimal medium supplemented with ferric citrate as indicated. The wash and the second elution fractions from each purification were loaded onto the SDS-PAA gel and analyzed by silver staining. The positions of the intrinsically biotinylated proteins PycA and AccB as well as of Fur-FLAG and Fpa-Strep are shown. The lower panel shows a Western blot using antibodies raised against the FLAG tag to detect the Fur-FLAG protein.

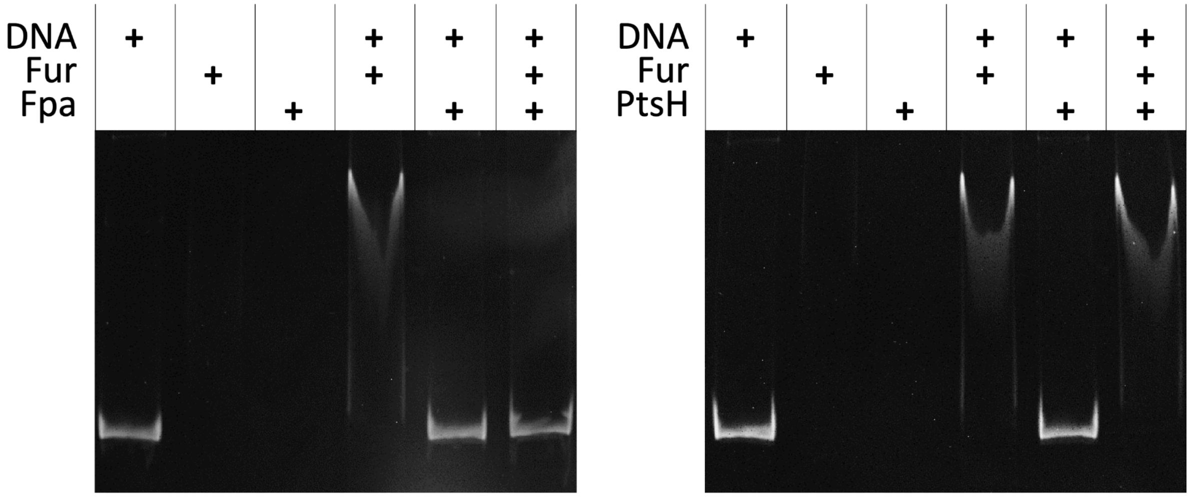


**Figure 5. Fpa is an antagonist of the DNA-binding activity of Fur.**
Gel electrophoretic mobility shift assay of Fur binding to *dhbA* promoter fragments. The components added to the assays are shown above the gels.
